# *TECPR2* a positive regulator of autophagy is implicated in healthy brain ageing

**DOI:** 10.1101/157693

**Authors:** John Alexander, Thomas Ströbel, Marianthi Georgitsi, Michael Schuster, Thomas Penz, Christoph Bock, Selma Hönigschnabl, Angelika Reiner, Peter Fischer, Peristera Paschou, Gabor G. Kovacs

## Abstract

Understanding the healthy brain aging process is key to uncovering the mechanisms leading to pathological age-related neurodegeneration, including progression to Alzheimer’s disease (AD). Here, we report the first deep whole genome sequencing study aiming to identify variants that are associated specifically to healthy brain aging defined on both clinical and neuropathological level, thus tacking the issue of pathological heterogeneity that often underlies a clinical AD diagnosis. We studied samples from the VITA brain bank and followed an extreme phenotypic ends study design comparing neuropathologically “healthy” aging individuals above 80 years of age with pure AD patients of the same age. Focusing on the extreme ends of the phenotypic distribution, and potentially functional variants, we discover a single variant *(rs10149146)* carried by 53.6% of the “healthy” brain elderly individuals in our study (15/28 individuals) and none of the 12 AD cases. This variant lies on the autophagy and cell cycle associated *TECPR2* gene. Autophagy dysfunction has been previously implicated in multiple progressive neurodegenerative diseases. An additional non-synonymous variant on the *CINP* gene (encoding a cell-cycle checkpoint protein) is also found in 46% of healthy controls and absent from all the AD cases. *TECPR2* and *CINP* appear to be “partner” genes in terms of regulation and their associated transcription factors have been previously implicated in AD and neurodegeneration. Our study is the first to support the hypothesis that a *TECPR2* non-synonymous variant carries a significant neuroprotective effect pointing to key molecules for the involvement of autophagy and cell cycle control in protection from neurodegeneration.

## INTRODUCTION

As life expectancy increases and fertility rates fall, the world population is aging^1^. However, despite the longer periods of good health and extended periods of social engagement and productivity, aging may also be associated with more illness and dependency. One of the consequences of longer life expectancies is the increase in people with dementia and especially Alzheimer’s disease (AD). The global projection of people living with dementia by 2050 is 115.4 millions^2^, posing a major burden on society and rendering imperative the need for understanding disease etiology and developing prevention strategies.

AD and other late onset forms of common neurodegenerative disorders are multifactorial and heterogeneous in nature and their genetic background remains poorly understood. Apart from their complex etiology (multiple genetic susceptibility factors), interaction with lifestyle and environmental factors leads to the onset of symptoms. Considering AD and including the *APOE* s4 haplotype, approximately 61% of the population attributable risk of late onset AD has been explained^3^. A recent meta-analysis of 74,046 individuals confirmed eight and reported 11 newly associated AD susceptibility loci^4-7^ A gene-based approach of the same dataset uncovered two more susceptibility loci^8^. However, the abovementioned genome-wide association studies implicate variants that account only for a small proportion of the estimated heritability of AD, leaving the substantial rest of the proportion unidentified. Attention has been turned to rare variants present in less than 1% of the population under the “common disease-multiple rare variants” hypothesis, currently amenable to large-scale analysis via next generation sequencing technologies^9^. In fact, low-frequency missense variants have been found to confer either strong protection or elevated risk of AD and cognitive decline^10-12^.

On the other hand, longevity and healthy aging show a heritability of 20-50%^13,14^. One might anticipate that genetic factors which increase the risk of common complex neurodegenerative diseases such as AD would negatively affect life-span and be less common among long-lived people as compared to younger aged individuals^15^. Conversely, the genetic contribution to longevity is hypothesized to be greatest at the oldest ages^16^; however, not much is known regarding how genetic factors might influence healthy brain aging and cognition maintenance^17^.The *TOMM40-APOE* region on chromosome 19q13.32 remains the most significantly associated locus with both longevity and cognition^16,18^.

To date, most genetic studies are based on a definition of case status using only clinical criteria. However, presence of multiple pathologies (i.e., combined neurodegenerative and vascular disorders) in the elderly with cognitive decline is rather the rule than the exception^19,20^. Therefore, pure AD pathology in demented individuals above 80 years of age is less frequent^19^. Moreover, individuals, including younger than 40 years of age, without clinical symptoms of dementia might show already AD-related changes in the brain even to a moderate degree (e.g., Braak stage III-IV of neurofibrillary degeneration^21^ or early phases of A-beta deposition^22^). Neurofibrillary tau pathology can appear without A-beta deposition called primary age-related tauopathy (PART). The present study is the first to attempt a genome-wide analysis aiming to identify variants that are associated to healthy brain aging defined on both clinical and a neuropathological level. In an effort to characterise factors protecting against pathological age-related neurodegeneration, we implemented an approach that focuses on the extreme ends of the phenotypic distribution under study (i.e. a comparison of normal aging individuals >80 years versus individuals with pure AD, lacking other proteinopathies or vascular lesions, of the same age). In a community-based study involving demented and non-demented individuals each represented approximately 6-10% of the cohort^19^, understanding such protective genetic factors may influence the adoption of strategies geared to prevent or decrease risk events that accelerate neurodegenerative changes in the susceptible aging brain when the processes are still reversible.

## RESULTS

Following our alignment and quality controls steps, deep whole genome sequencing resulted in 10,908,452 variants in total, including 40,435 nonsynonymous, 475 stop-gain and 39 stop-loss variants annotated by RefSeq database (Table 1). Our analysis of variants previously implicated in AD replicated the positive association to ApoE allele (p-value 4.37e-05 (33% in Cases, 1.8% in Controls) and we were further able to pick up a neuroprotective effect for the ε2 allele (p-value of 0.02 - 0%Cases,17.9% Controls) (See Supplementary results). Looking into variants that have been previously implicated in longevity, we also picked up the association to *rs4420638* marker near *APOC1* gene which was previously implicated in a case-control GWAS conducted on 763 German centenarians and nonagenarians, and 1,085 controls (mean age 60 years)^16^ (P value =2.00e-04, see Supplementary results for details).

**Table 1.**
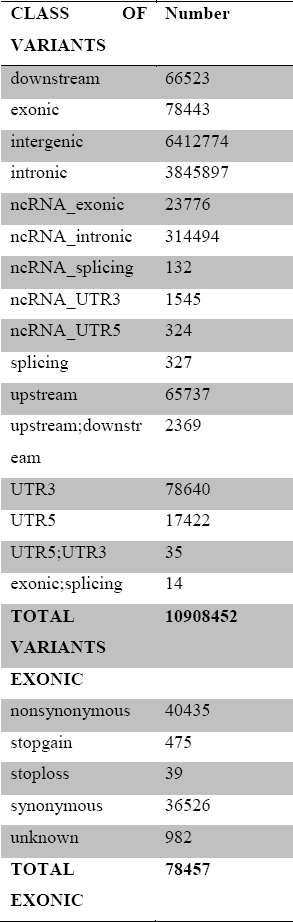
Summary statistics on whole genome sequencing data from 40 studied individuals after applying quality control steps. **ANNOVAR definitions** **Exonic** - variant overlaps a coding region; **Splicing** - variant is within 2-bp of a splicing junction; **ncRNA** - variant overlaps a transcript without coding annotation in the gene definition;**UTR5** - variant overlaps a 5' untranslated region;**UTR3**- variant overlaps a 3' untranslated region; **Intron** - variant overlaps an intron;**Upstream**- variant overlaps 1-kb region upstream of transcription start site;**Downstream** - variant overlaps 1-kb region downtream of transcription end site;**Intergenic** - variant is in intergenic region

To examine variants most likely to have a neuroprotective effect, we filtered for unique variants in at least 10 of the 28 neuropathologically “normal” controls, in our study but absent in all 12 cases. This resulted in 6,984 variants (Table S1), including 55 non-synonymous and one stopgain variant. Further removal of sequence artefacts and pseudogenes resulted in 14 prioritized exonic variants (Table 1, S2). A summary of these genes and their functions are provided in Table 2. Besides gene-coding variants, we also explored the potential involvement of variants in exons of non-coding RNAs. Again, as a filtering step, we focused on variants that are common in at least 10 “healthy” aged controls and absent from all cases. We identified eight such prioritized variants as shown in Table 3.

**Table 2.**
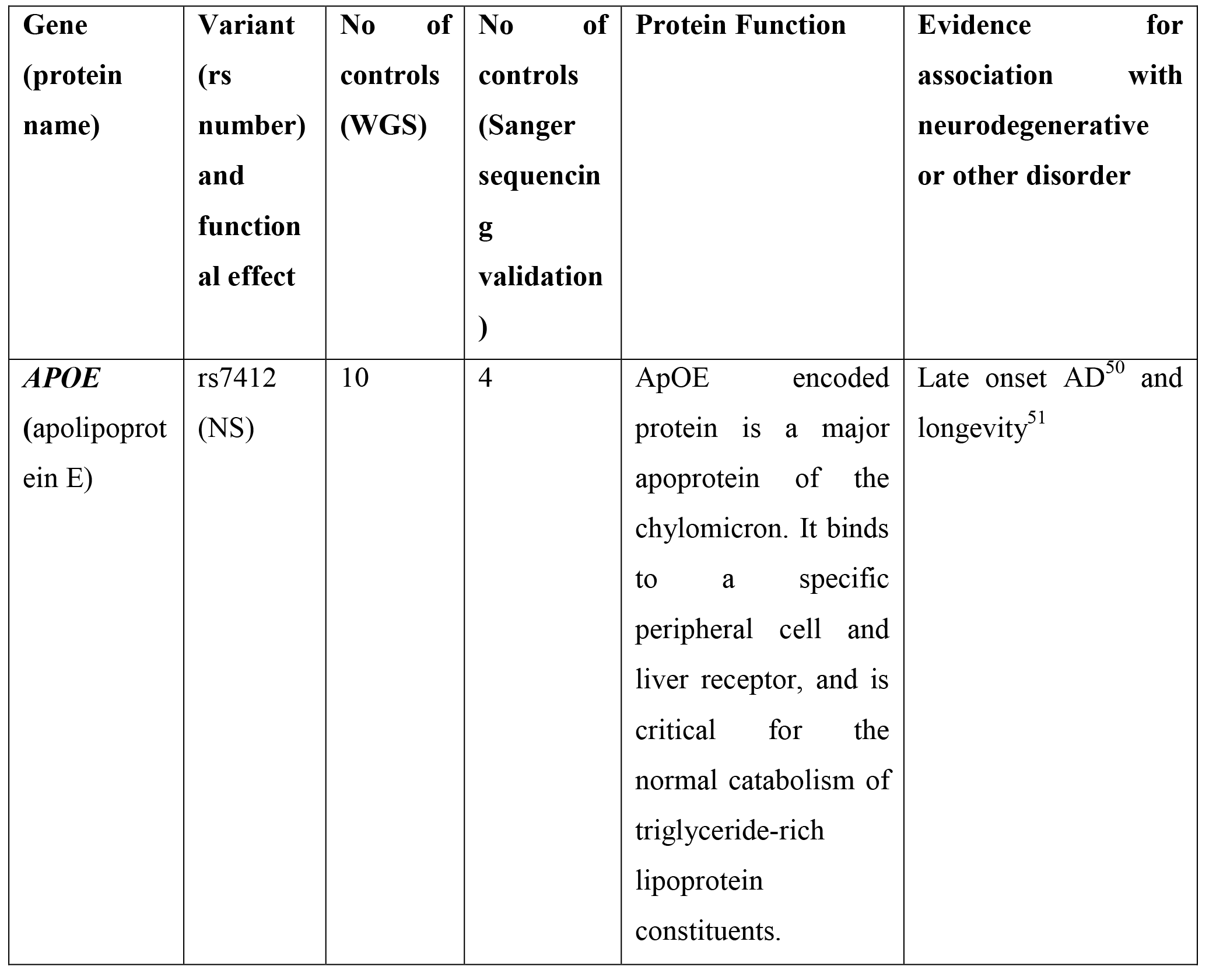
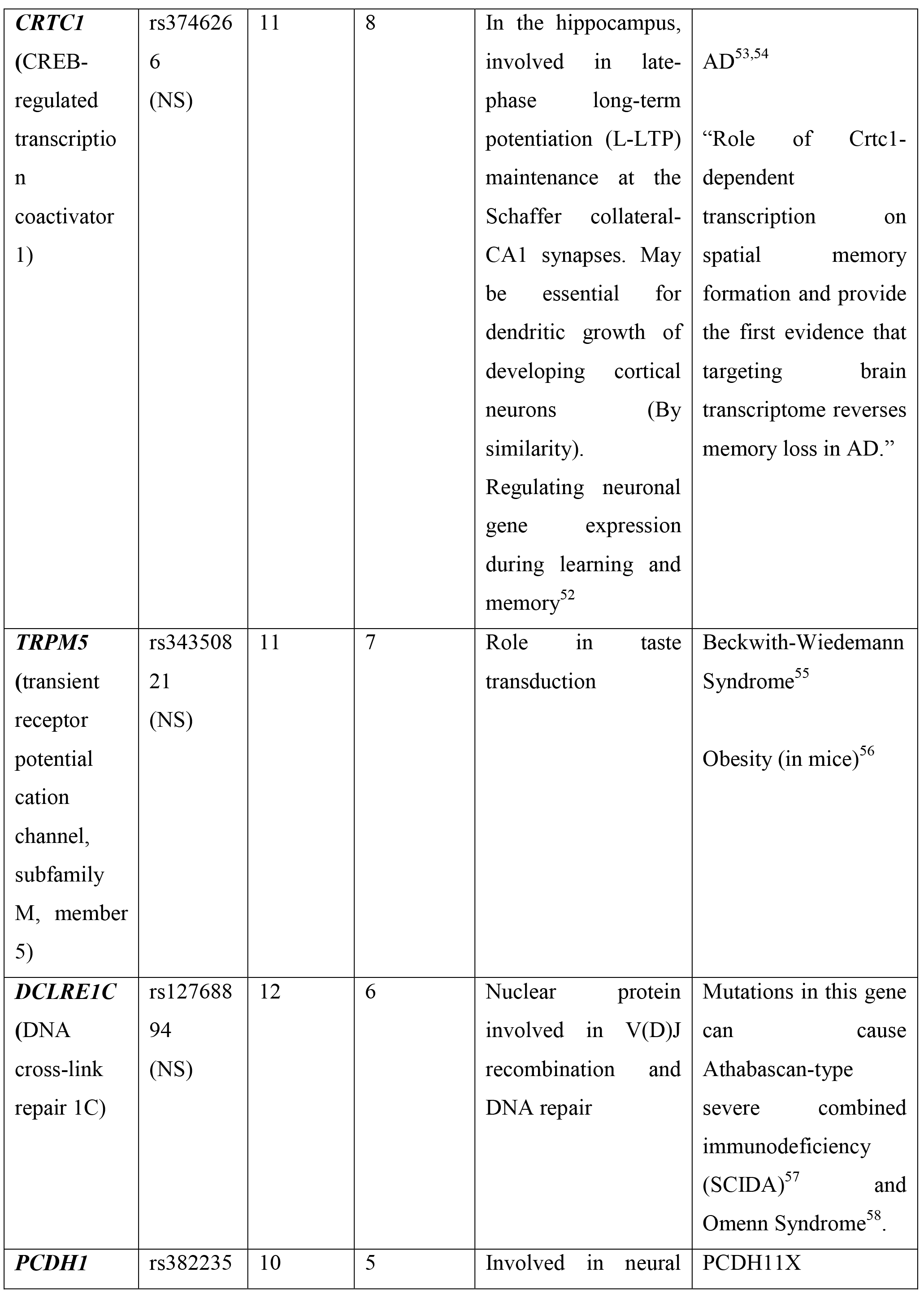
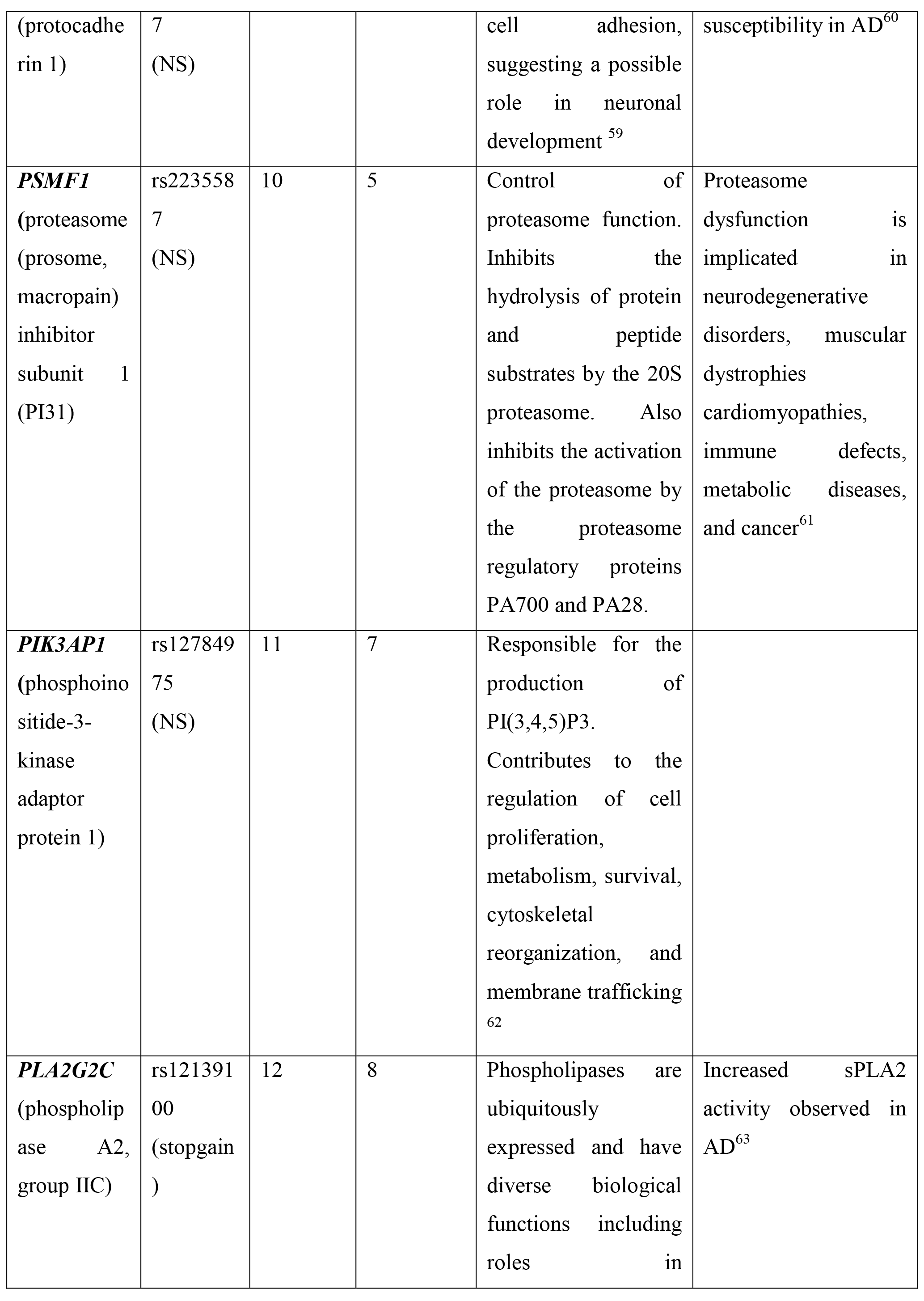
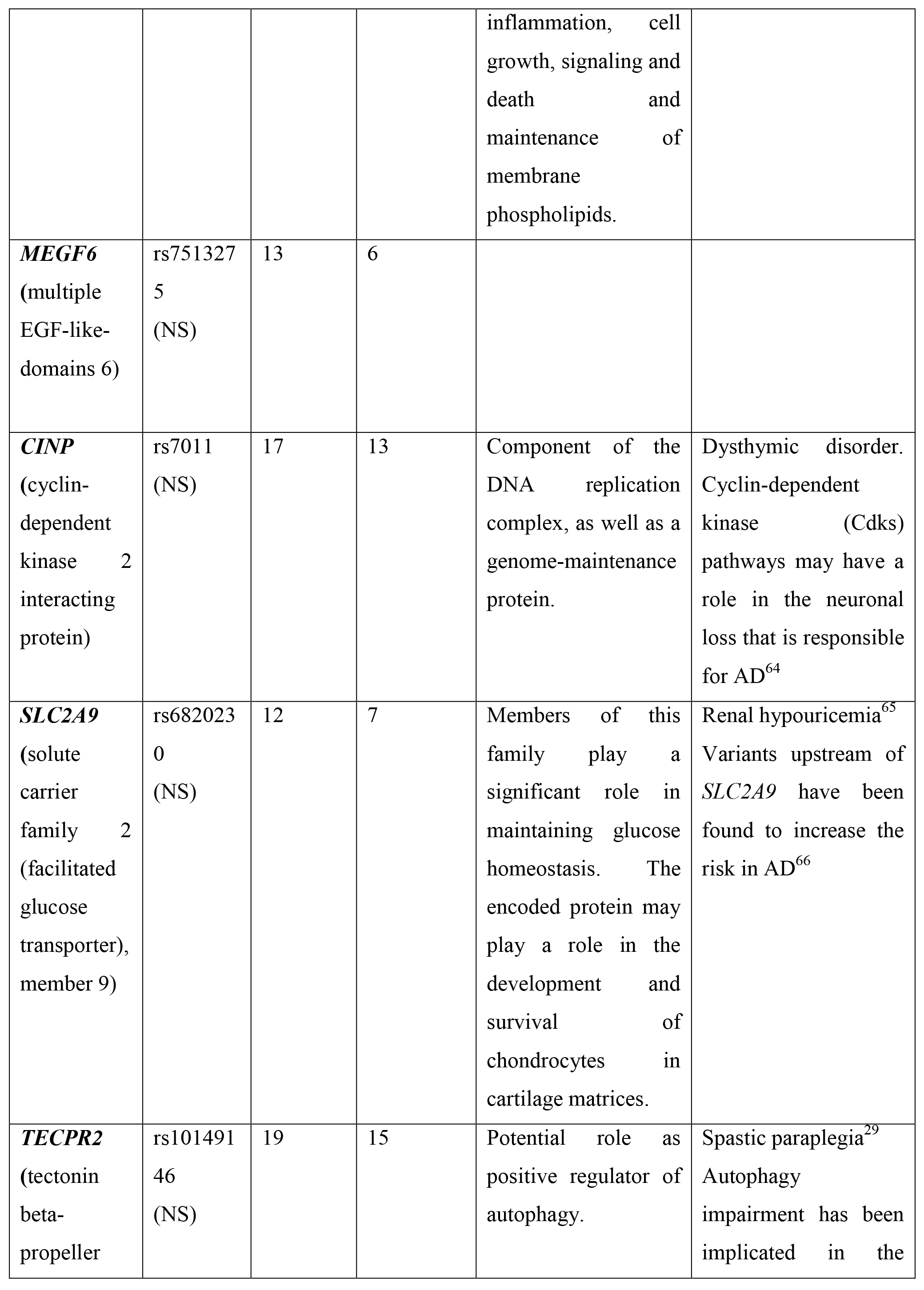
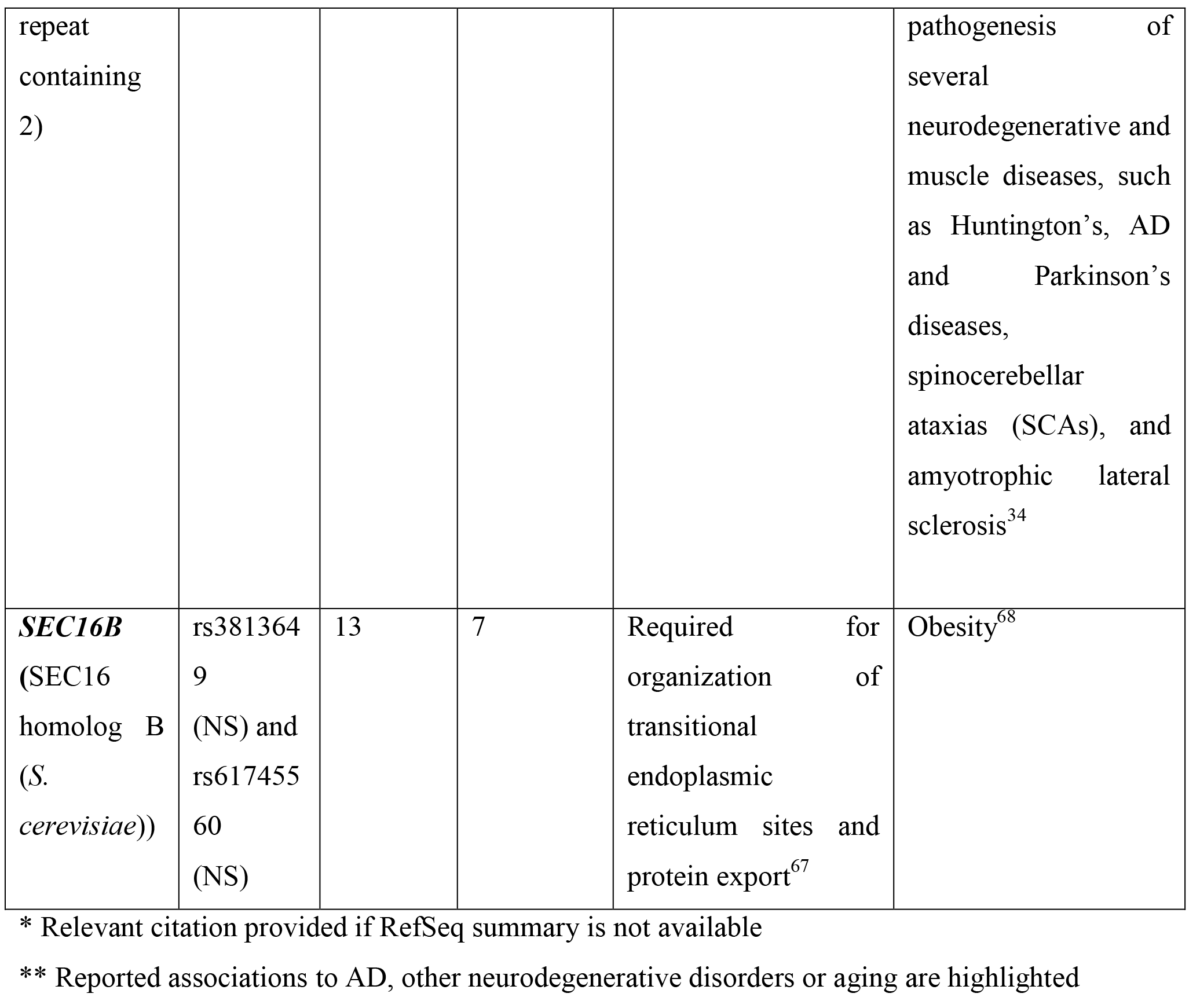
Unique exonic variants (NS-nonsynonymous, stopgain) residing in protein-coding genes and present in at least 10 controls and absent from all 12 AD cases following whole genome-sequencing (WGS) analysis. Sanger sequencing validations are also shown..

**Table 3.**
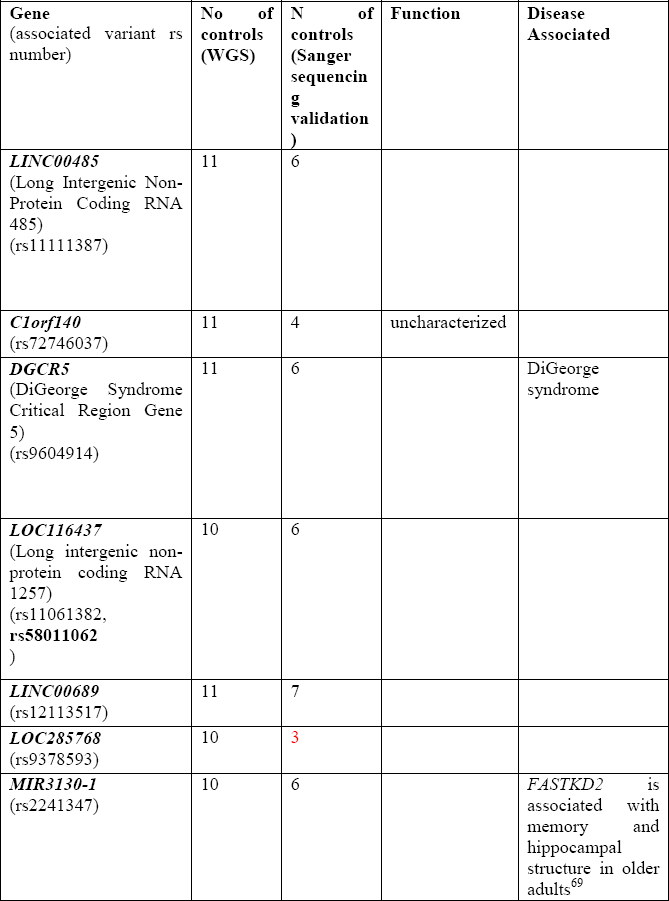
Unique variants identified in exons of ncRNA genes in at least 10 controls and absent from all 12 AD cases following whole genome sequencing (WGS) analysis. Sanger sequencing validations are also shown.

We proceeded to perform Sanger sequencing in order to confirm the whole genome sequencing results. Sanger sequencing was not able to confirm all of the whole genome sequencing called variants. We were thus able to further narrow down our search for genes that carry non-synonymous variants in at least 10 controls and none of the cases to only two genes; The Tectonin beta-propeller repeat containing 2 gene *(TECPR2),* with allele G on *rs10149146* present in 15 controls and no AD cases and the Cyclin-dependent kinase 2 interacting protein *(CINP)* with allele T on *rs7011* present in 13 controls and absent from all cases. The Fisher’s exact test revealed statistically significant association to healthy brain aging with SNP *rs10149146* of *TECPR2* (P=4.055e-4). Interestingly, the two aforementioned SNPs lie physically close (86.16 kb) and are in high linkage disequilibrium (r^2^ =0.73). The genes carrying these variants have been previously found to share a bidirectional promoter thus being subject to common gene expression regulation^23^. Our analysis showed that transcription factors that potentially co-regulate the two genes *(TECPR2* and *CINP)* include OCT1 (Octamer-Binding Transcription Factor 1 - POU2F1), NMYC (N-myc proto-oncogene protein), USF (upstream transcription factor 1), TAX/CREB (Tax/cAMP response element-binding protein), HEN1 (Helix-Loop-Helix Protein 1), YY1 (Yin And Yang 1), and RFX1 (Regulatory Factor X1).

## DISCUSSION

In the past two decades, evidence has accumulated in favour of the existence of genetic factors associated with AD-related pathology, via either GWAS studies with single-marker associations alone or by combining GWAS data with neuroimaging data, leading to the identification of 20 AD susceptibility loci^4-8,24-26^. However, very little is known about the protective genetic influences against neurodegeneration and in favour of healthy brain aging. Genetic influences of cognitive abilities and aging in old age are prominent and approximately 50% of this variance may be attributed to multiple genetic variants with small effects^27^. Candidate gene studies have highlighted the potential roles of genes such as *BDNF, COMT* and *DTNBP1* in either cognitive ability and normal function (such as learning and memory) or cognitive decline; yet associations for other genes remain inconclusive^28^.

In this study, we applied a genome-wide sequencing analysis aiming at the identification of variants and genes that may be important for healthy aging, in an effort to characterise factors for the prevention of pathological age-related neurodegeneration. Importantly, based on detailed neuropathological evaluations, we were able to define two extremes of the phenotypic distribution of brain aging; old age individuals with minimal signs of neurodegeneration versus pure AD. Demonstrating the power of our approach, despite the small sample size, we were able to pick up the neuroprotective effect of the *APOE2* variant (p=0.02). We identified two non-synonymous exonic variants in two genes that are present in 53.6% ad 46% respectively of 28 healthy aging controls and absent from all 12 pure AD cases that we studied. The associated genes *TECPR2* and *CINP* appear to be “partner” genes in terms of regulation^23^ and are associated with autophagy^29^ and cell-cycle control^30^ respectively. Notably, the transcription factors we found associated with these two genes have also previously been implicated in neurodegeneration; a *POU2F1 (OCT1)* intronic variant has been associated with AD and the *POU2F1* gene was found to be down-regulated in the hippocampus of AD brains but not healthy controls^31^, USF has been found to interact with the Alzheimer amyloid beta-protein precursor gene^32^, The beta-amyloid peptide mediates synapse loss through the CREB signaling pathway, while YY1 is an activator of beta-site amyloid precursor protein-cleaving enzyme 1 (BACE1), a prerequisite for the generation of beta-amyloid peptides, the principle constituents of senile plaques in the brains of patients with Alzheimer’s disease (AD)^33^..

A variant in *TECPR2* (namely *rs10149146,* c.A2047G, p.I683V) was present in the highest number of our control individuals (n=15/28). *TECPR2* has been reported to be a positive regulator of autophagy and involved in hereditary spastic paraparesis^29^. Autophagy dysfunction is implicated in multiple progressive neurodegenerative diseases, and has been reported to play a major role in the pathogenesis of several neurodegenerative and muscular diseases, such as Huntington’s Disease, AD and Parkinson’s Disease, spinocerebellar ataxias, and amyotrophic lateral sclerosis^34^. Mutations in *TECPR2* have been also recently found to cause a subtype of familial dysautonomia comparable to hereditary sensory autonomic neuropathy with intellectual disability. *TECPR2* is also a possible candidate gene for human neuroaxonal dystrophies (NAD), which may be associated with aging and supplementary to numerous metabolic-toxic conditions^36^.

On the other hand, 13 controls were found to share a variant in *CINP* (rs7011, aNP:NM_001177611:exon5:c.G536A:p.R179H,

CINP:NM_032630:exon5:c.G491A:p.R164H) which was however absent from all of our AD cases. The protein encoded by *CINP* is a member of the cyclin-dependent kinase (CDK) family of Ser/Thr protein kinases and has been identified as a genomic maintenance protein^30^. It is a cell-cycle checkpoint protein, interacting with ATR-interacting protein to regulate ATR-dependent signaling, resistance to replication stress, and G2 checkpoint integrity^30^. Cdk2 is also a key regulator of the senescence control function of Myc and a potential therapeutic target for treating tumors derived by Myc or Ras ^37^. Deregulation of protooncogenes like MYC and RAS augment tumor development although normally they function to promote normal cell growth and may have an impact on tissue regeneration^37^. An increasing body of studies show that cell cycle disturbances may play an early role in AD pathogenesis^38-40^. CDKs^41^ propel tau phosphorylation. *In vitro* assays have indicated that ectopically re-expressed CDKs in neurons in AD phosphorylate tau in a similar manner to that found in AD *in vivo^41^*. Increased N-MYC and C-MYC expression in reactive astrocytes may also play a role in reactive astrocytosis in human neurodegenerative disorders^42^.

Despite the relatively low number of cases examined, our strict criteria and by focusing on extreme ends of the phenotypic distribution of neuropathology-related variants, we were able to identify an autophagy and cell-cycle related gene to be associated with healthy aging specifically lacking pathological protein depositions and vascular lesions, which is unusual in the aging brain^19,20^. The implicated genes warrant further investigation and highlight the key role of autophagy and cell-cycle control in protecting from neurodegeneration. This is the first study to specifically attempt to find genetic variants that promote healthy brain aging based on neuropathological evaluations of post-mortem samples. Our approach contrasts those based only on clinical criteria but also neuropathology-based studies, which do not apply a holistic approach by examining the whole spectrum of proteinopathies and other brain lesions. In an aging world and with no existing cure for dementias, the need to increase our understanding of their etiology is imperative, Elucidating the physiological pathways that drive the shift from healthy brain aging to neurodegeneration, will lead to the identification of targets for improved therapies and prevention strategies for AD and related neurodegenerative disorders.

## METHODS

### Subjects

We selected cases from the longitudinal aging transdanubian VITA study^19^, which included individuals after written consent of either the patient or the family or legal representative of the patient. The Ethical Committee of the Medical University of Vienna gave approval for the genomic analysis (Nr: 140/2011). All experiments were conducted according to the principles expressed in the Declaration of Helsinki, and in accordance with relevant Austrian guidelines and regulations, including the right to object from participating in scientific research. The manuscript does not contain information or images that could lead to an identification of the individual or which could violate any personal rights. Bioinformatic processing was performed anonymously. Twelve participants with pure end-stage AD with only beta amyloid plaques (Thal phase 5) and neurofibrillary tangles (NFTs; Braak stage VI) in the brain and no other comorbidities (Lewy body pathology, TDP-43 proteinopathy, other tauopathies, vascular diseases etc.) participated (for details on neuropathological characterisation see^19^). A typically aging group of 28 participants aged 80 and above with minor amount of NFTs (Braak stage I but lack of A-beta plaques), not enough to be suggestive of AD and with no other neurodegenerative or vascular disorders were used as a control group.

### Whole genome sequencing (WGS), processing of raw sequences and variant calling

DNA was extracted from fresh-frozen cerebellum and frontal cortex tissue. Whole genome sequencing was performed by the Biomedical Sequencing Facility at CeMM, using Illumina HiSeq 2000 machines with a read length was 2x100 base pairs in the paired-end configuration. The sequencing reads were aligned to the b37 reference genome with Epstein-Barr Virus sequence (BROAD 1000 Genomes GRCH37) using BWA-MEM aligner, and variant calling was done following the GATK^43^ pipeline from GATK Best Practices available at the time of processing (http://www.broadinstitute.org/gatk/). Each sample was preprocessed to remove PCR and optical duplicates using Picard. In order to improve the SNP calling, INDEL realignment was done using IndelRealigner and recalibration of base quality scores was performed using baseRecalibrator from Genome Analyzer Toolkit, to eliminate systematic errors per lane. Once recalibrated, variant calling and recalibration was performed on the merged samples using GATK version 3.1-1 Haplotype Caller. All samples achieved the goal of 30x genome-wide coverage with a mean coverage of 37.23x (4.47 Terabases of data). Only variants that passed the quality control filters (quality score QUAL>=30) and genotype threshold of 95% were kept for further analysis. Additional quality control steps were performed using vcftools (version 0.1.9)^44^ and statistical analysis was done in R (version 3.0.1). *Variant Ranker* (http://paschou-lab.mbg.duth.gr/Software.html) was used to annotate and filter the variants. *Variant Ranker’s* CaseControl filtering module utilizes SnpSift tool (CaseControl)^45^ to count the number of homozygous, heterozygous and total variants in the case-control groups and implements ANNOVAR ^46^ for annotation of variants. Haploview^47^ was used to perform Fischer’s exact test.

### Filtering of Variants and Case-Control analysis

RefSeq was used as the reference database for gene definitions. Our analysis primarily focused on functionally important exonic variants (i.e. non-synonymous/stopgain/stoploss/splicing effect) and variants present in exons of non-coding RNA genes (ncRNAs). In order to examine unique variants/mutations only present in the control samples with a likely neuroprotective effect, we filtered for variants that were carried by at least 10 control individuals (i.e. control subjects carried at least one alternative allele) and were absent in all 12 cases (i.e. cases carried only the reference alleles for the same locus in the human reference genome). Sequence artefacts and pseudogenes were removed from our analysis by examining gene exclusion lists in *Fuentes Fajardo 2012^48^.* Database for Annotation, Visualization and Integrated Discovery (DAVID)^49^ was used for gene ontology analysis.

### Sanger Sequencing

All controls carrying prioritized variants were sequenced by the Sanger custom sequencing method, using the BigDye Terminator v.3.1 chemistry on an ABI 3730XL genetic analyzer. Sequencing was performed on both DNA strands and sequences were both aligned and visually observed by at least two independent researchers.

## ACKNOWLEDGEMENTS

This study was performed in the frame of the EU FP7 Project DEVELAGE (#278486; G.G.K.).

## Author contributions.

JA performed the statistical analysis, interpreted findings, and wrote the manuscript; PP and GGK were responsible for study design, supervised analysis and interpreted findings and wrote the manuscript; GGK supervised neuropathological evaluations; TS processed samples and performed sanger sequencing analysis; MG participated in the interpretation of results and wrote the manuscript; MC, TP, and CB performed whole genome sequencing analysis; SH, AR, PF, were responsible for the VITA study including clinical, pathological and neuropathological evaluations; all authors contributed to writing and revision of the manuscript

